# mNGS Investigation of Single Ixodes pacificus Ticks Reveals Diverse Microbes, Viruses, and a Novel mRNA-like Endogenous Viral Elements

**DOI:** 10.1101/2022.08.17.504163

**Authors:** Calla Martyn, Beth M. Hayes, Domokos Lauko, Edward Mithun, Gloria Castañeda, Angela Bosco-Lauth, Amy Kistler, Katherine S. Pollard, Seemay Chou

**Affiliations:** Department of Biochemistry & Biophysics, University of California – San Francisco, San Francisco, CA; Gladstone Institute of Data Science & Biotechnology, San Francisco, CA 94158; One Health Institute, Colorado State University - Fort Collins, CO 80523; Department of Biomedical Sciences, Colorado State University - Fort Collins, CO 80523; Chan Zuckerberg Biohub, San Francisco, CA 94158; Department of Epidemiology & Biostatistics, University of California - San Francisco, San Francisco, CA 94158

## Abstract

Ticks are increasingly important vectors of human and agricultural diseases. While many studies have focused on tick-borne bacteria, far less is known about tick-associated viruses and their roles in public health or tick physiology. To address this, we investigated patterns of bacterial and viral communities across two field populations of western black-legged ticks (*Ixodes pacificus*). Through metatranscriptomic analysis of 100 individual ticks, we quantified taxon prevalence, abundance, and co-occurrence with other members of the tick microbiome. Our analysis revealed 11 novel RNA viruses from *Rhabdoviridae, Chuviridae, Picornaviridae, Phenuiviridae, Reoviridae, Solemovidiae, Narnaviridae*, and 2 highly divergent RNA viruses lacking sequence similarity to known viral families. The majority of these viruses were also detectable in lab-raised ticks at all developmental life stages, localize to tick salivary glands, and show evidence of circulation in mice fed on by ticks. These data suggest that viruses are stable, heritable, and transmissible members of the tick microbiota. We also unexpectedly identified numerous virus-like transcripts that are associated with tick genomic DNA, most of which are distinct from known endogenous viral element-mediated immunity pathways in invertebrates. Together, our work reveals that in addition to potentially serving as vectors for potential viral pathogens, *I. pacificus* ticks may also have symbiotic partnerships with their own vertically-transmitted viruses or with ancient viruses through evolutionarily acquired virus-like transcripts. Our findings highlight how pervasive and intimate tick–virus interactions are, with major implications for both the fundamental physiology and vector biology of *I. pacificus* ticks.

## Introduction

Ticks are increasingly important disease vectors for humans and livestock, particularly in the United States, where they account for more cases of vector-borne diseases than mosquitoes. Approximately fifty thousand confirmed cases of tick-borne diseases are reported annually^1^, which is likely an underestimate due to diagnostic challenges associated with Lyme disease and our poor understanding of rare tick-borne diseases or diseases of unknown etiology. Currently, the majority of field surveillance studies of tick-associated microbes focus on the causative agent of Lyme disease *Borrelia burgdorferi* and a select number of other known human pathogens, such as *Rickettsia* (Rocky Mountain spotted fever and other Rickettsioses), *Anaplasma phagocytophilium* (Anaplasmosis), and Powassan virus. Although the full diversity of microbes carried by ticks is much greater than those definitively linked to human disease^2–21^, we know strikingly little about the ecology or disease implications of most tick-associated microbes.

Furthermore, we are only beginning to appreciate the broader role microbes play in tick biology. Like many other invertebrates, tick–microbe interactions go far beyond the transmission of human pathogens. Tick–microbe interactions can be antagonistic, neutral, or beneficial^22–25^. Some microbes, such as the bacterial endosymbiont *Rickettsia*, play fundamental roles in tick physiology through stable and symbiotic interactions^25^. It is not as well known whether ticks also form stable, symbiotic interactions with viruses and how such interactions may also influence tick biology. Some viruses have been identified not only in live ticks but also in many laboratory-passaged tick cell lines^26^. Microbes have also been implicated in shaping the evolution of ticks through horizontal transfer of bacterial genes^27^ and endogenization of viral sequences as an immune response^28–31^.

Although the relationship between ticks and viruses is a subject of much interest and field microbiome studies have increased our catalog of tick-associated microbes^32^, there still remain several outstanding questions. A number of experimental strategies and technical hurdles have limited the scope and depth of microbiome analyses in ticks. First, field studies often sequence pooled tick samples, preventing quantitative examination of microbial prevalence, co-occurrence, and per-sample relative abundance. These metrics could greatly enable more sophisticated analyses of transmission dynamics and ecology. Furthermore, our ability to capture lower abundance microbes is hampered by the dominance of tick host sequences in genomic and metagenomic libraries. This limits our understanding to the most abundant microbes in ticks, which does not necessarily coincide with all microbes that have important roles in disease or tick physiology.

Finally, our understanding of tick microbiota in North America is currently biased towards species historically associated with human diseases, such as *Ixodes scapularis*, the primary vector for Lyme disease in the Eastern United States. In recent years, there has been an expansion of tick-borne disease cases on the West coast of the U.S. that have been attributed to other less-studied tick vector species. For example, *Ixodes pacificus* is a major tick species extending from Northern Mexico to British Columbia^33^. *I. pacificus* ticks are most abundant in California, where they cover 96% of all counties^34^ and are responsible for the majority of human tick bites^35^. They are vectors for a variety of well-characterized human pathogens such as *B. burgdorferi, Borrelia miyamotoi, Babesia odocoilei, Bartonella spp, A. phagocytophilum*, and *Ehrlichia spp*^36^. Despite this, *I. pacificus* is substantially understudied compared to the eastern black-legged tick *I. scapularis.* For these reasons, we chose to focus this study on *I. pacificus* ticks.

To provide much-needed insight into tick-borne microbes in the Western U.S., we examined the microbiomes of *I. pacificus* ticks collected from two coastal habitats in California where humans are likely to encounter them^35,37,38^. In order to capture lower-abundance microbes, we coupled an experimental microbial enrichment workflow with RNA sequencing to profile both bacteria and RNA viruses. Analysis of microbiomes at the level of individual ticks enabled us to also quantify patterns of microbial prevalence and abundance. We performed follow up laboratory-controlled experiments examining microbial localization in both ticks and mouse bloodmeal hosts, which provided additional insights into potential transmission dynamics and the symbiotic nature of tick-virus relationships.

## Results

### Establishment of an RNA-based approach to defining composition of field tick microbiomes

We set out to define the metatranscriptome of *I. pacificus* ticks collected from coastal California, focusing on two sites associated with human exposure^35–39^. We examined the two most developmentally advanced life stages (nymphal and adult) that are more amenable to single-tick sequencing due to greater individual biomass. These sample sets included adult ticks from Garrapata State Park and nymphal ticks from China Camp State Park. Adults were collected in the Fall (2019) and nymphs were collected in the Spring (2020) so that we could investigate seasons and life stages enriched for human contact. In total, RNA libraries were sequenced for 100 individual ticks.

The majority of whole-tick RNA libraries are composed of tick ribosomal RNA, which reduces the power to detect lower abundance bacterial and viral sequences. To address this challenge, we enriched microbial sequences by experimentally depleting abundant tick sequences through Depletion of Abundant Sequences by Hybridization (DASH) (Figure 1a)^40^. For adult tick libraries, DASH-based depletion enriched non-host reads by nearly ten-fold (Figure 1b). Sequencing libraries generated for smaller nymphal ticks were not of sufficient concentration to effectively perform DASH. After quality filtering and host subtraction, non-host reads were first classified using the metagenomic classifier kraken2^41^ (Figure 1a). Samples varied substantially in the proportion of reads able to be classified, with anywhere from 0.2% to 83% of reads being classified by the tool (Figure 1c). All classified reads by kraken2 were bacterial; no known viruses were identified. Because of this, viruses were classified in a custom pipeline that enabled the detection of divergent viruses (Figure 3a, Methods).

**Figure 1:**
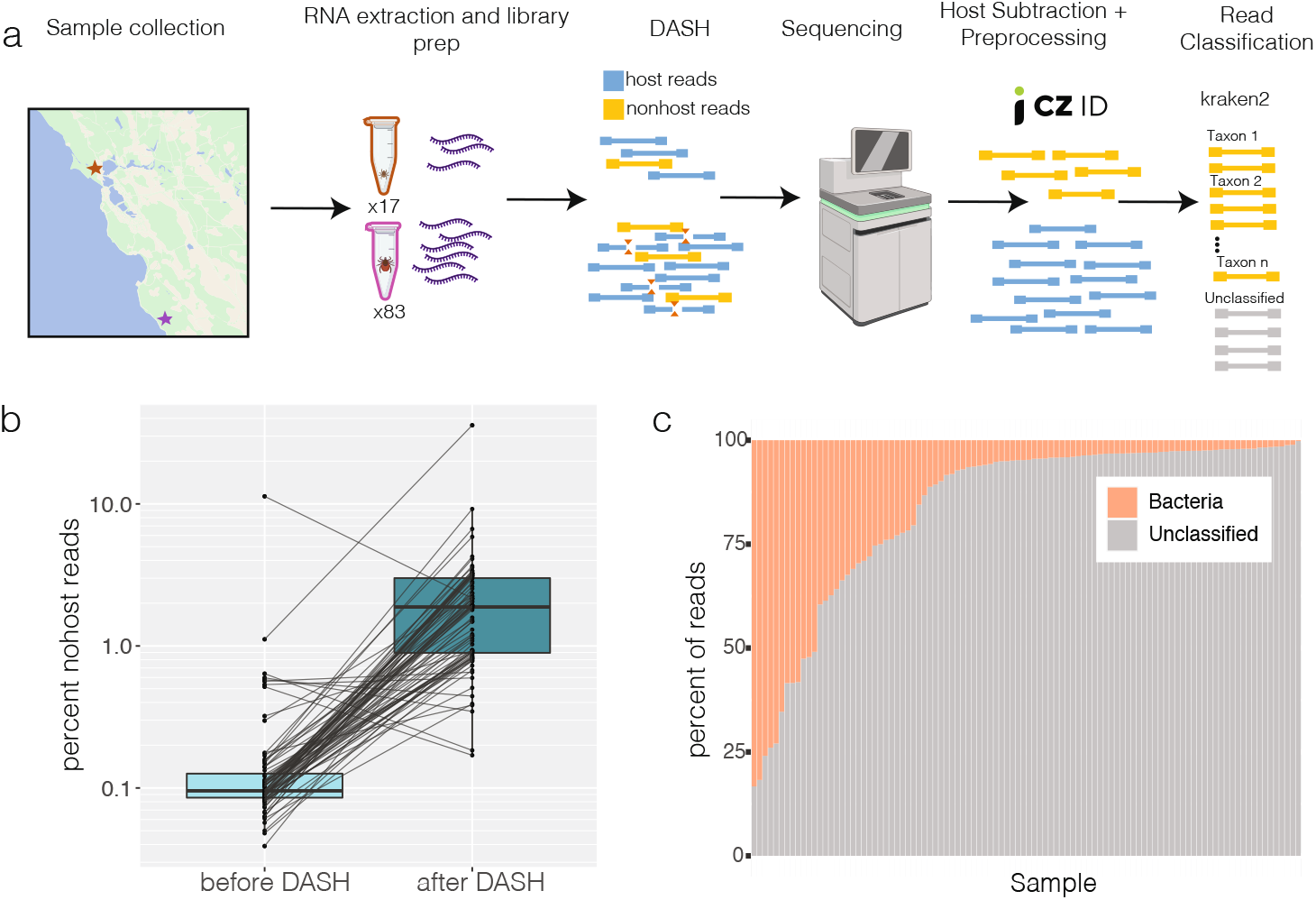
Experimental approach. a) 17 nymphs and 83 adults were collected from Garrapata State Park (purple) and China Camp State Park (red) respectively. RNA was extracted from whole bodies of individual ticks, mNGS librariers were prepared and DASH was performed to deplete abundant tick sequences. After sequencing, reads were quality filtered and tick-derived reads were removed using CZID. Remaining reads were classified by kraken2. b) The percent of non-host reads as classified by CZID for matched libraries before and after DASH, line connect individual libraries. c) Percentage of nonhost reads classifed per sample by kraken2.

To assess the general validity of our approach to characterizing the microbiota of field ticks, we first quantified the bacterial component of tick metatranscriptomes. We identified 114 bacterial genera across the dataset with a median of 11 genera per tick (Figure S1a). Larger libraries had more classified genera, indicating that sequencing depth is a limiting factor in characterizing tick microbial diversity (Figure S1a, S2). We compared the taxonomic composition of our samples with previously reported tick-associated human pathogens, such as *Rickettsia, Anaplasma*, and *Coxiella*, which are most commonly linked to *I. pacificus* ticks^3,6,15,36^. *Borrelia, Borreliella, Ehrlichia*, and *Bartonella* are also human pathogens known to circulate in this species, although typically at lower frequencies. *Ehrlichia, Borrelia*, and *Borreliella* were identified at rates of 2–3% across the full dataset but not in samples with a more stringent quality cut-off of at least 1 million non-host reads (Figure 2, Figure S2). We did not identify *Coxiella, Bartonella*, or *Francisella* in any ticks, indicating they are either absent in this population or present at levels too low to be detected.

**Figure 2:**
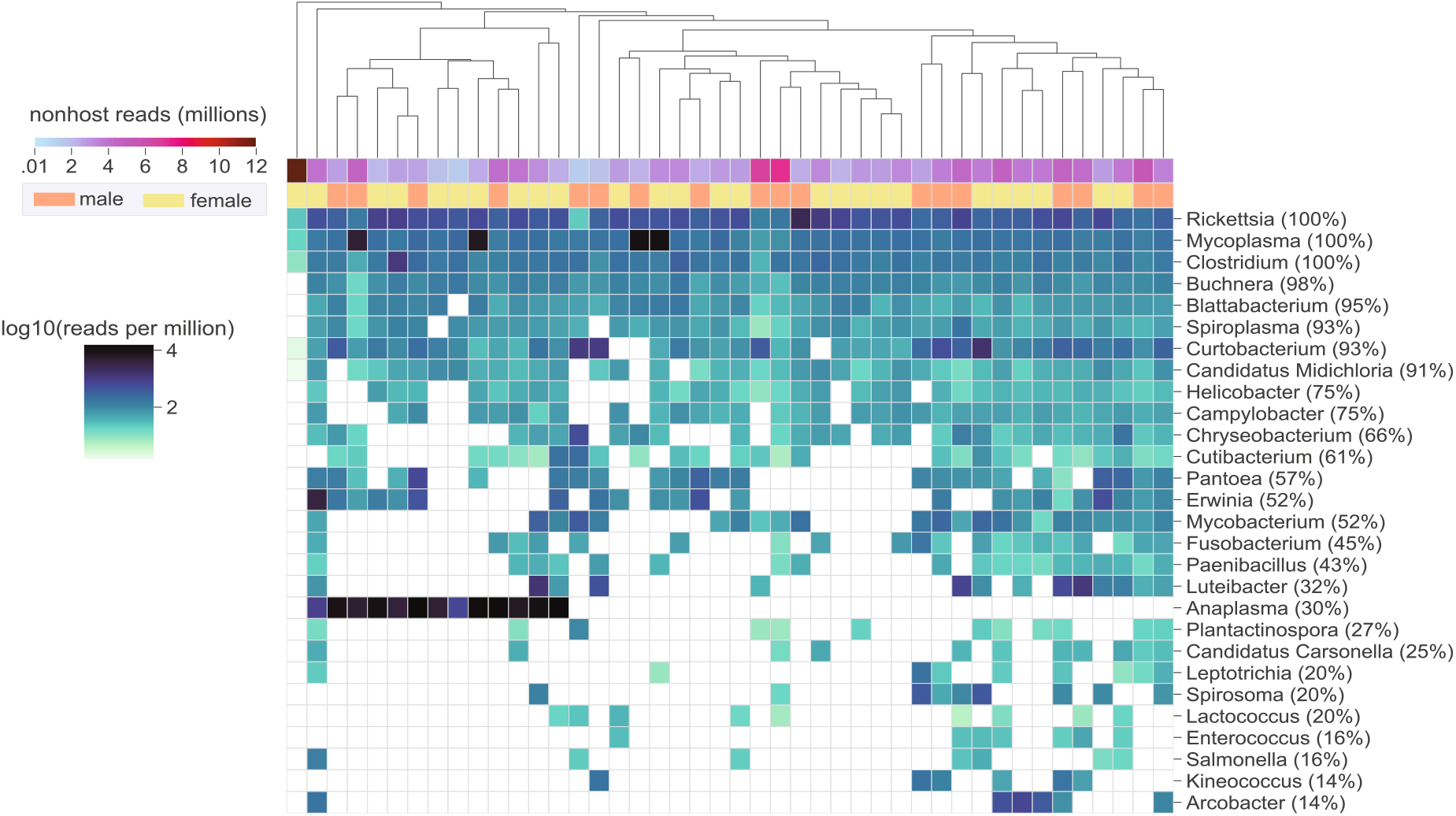
Bacterial genera detected in *Ixodes pacificus.* Heatmap displaying reads per million (rpm) of bacterial genera as classified by kraken2. Plot is limited to samples with at least 1 million nonhost reads and genera detected in at least 5 samples. Prevalence in the selected samples is shown next the genus name. Rows are ordered by decreasing prevalence and columns are heirarchically clustered by euclidean distance.

**Figure 3:**
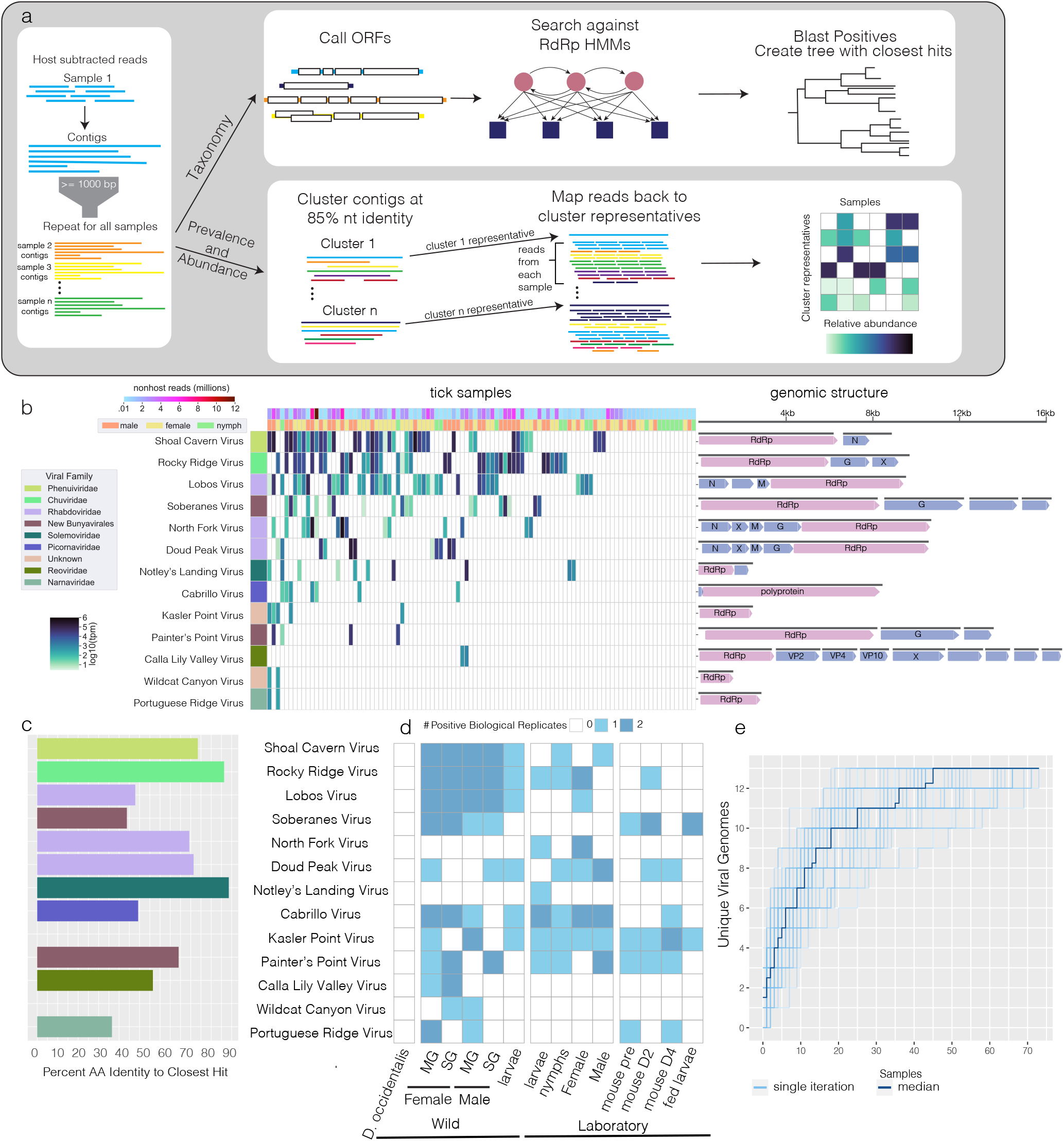
DIscovery of novel viruses in *Ixodes pacificus*. a) Analysis pipeline for identification of viruses: reads were assembled into contigs and open reading frames were predicted. Resulting proteins were scanned for the presence of an RdRp using HMMER and classified by viral family using the closest hits by blast. Prevalence and abundace was determined by clustering contigs and mapping reads back to cluster representatives. b) Heat map of transcripts per million (tpm) of each identified RdRp containing viral contig across the dataset, open reading frames are displayed to the right and annotated with identified protein (RdRP=RNA-dependent RNA polymerase, G=Glycoprotein, N=Nucleoprotein, M=Matrix protein, VP=Viral Protein). Open reading frames with “X” contain homology to known viral proteins of unknown function and open reading frames with no annotation had no identfieid homology to known proteins. c) Percent amino acid identify of each virus to nearest hit in blast, viruses with no bar had no blast hits. d) Table summarizing detection of each viral contig in tick and mouse samples. e) Rarefaction plot showing increase in number of viral genomes for each sample analyzed, samples were randomly shuffled 50 times, median shown in dark blue

Of the samples with at least 1 million non-host reads, several genera we identified were found at frequencies similar to previous reports. We found the endosymbiont *Rickettsia* in 100% and *Anaplasma* in approximately 30% of ticks (Figure 2). We also observed several instances of coinfections between *Ehrlichia, Borrelia, Borreliella*, and *Anaplasma* (Figure S1b). In total, results from our bacterial analyses are largely consistent with previously reported analyses of bacterial constituents of tick microbiota, suggesting our RNA-based approach to characterizing tick-associated microbes can indeed be applied to field studies. These results gave us confidence that our sequencing and analysis pipeline could reliably report on microbiomes of field *I. pacificus* ticks at nymphal and adult life stages.

### Optimized workflow enables detection of low-abundance bacteria and RNA viruses

Our combined approach of host depletion and RNA-sequencing opened up several unique lines of inquiry. In addition to known tick-borne human pathogens, we identified several bacterial genera previously undetected in *I. pacificus*. Although these genera had high prevalence across our samples, they were present at low relative abundances and likely escaped detection in previous studies in the absence of DASH-based microbial enrichment. *Mycoplasma*, a genus that has been linked to Lyme-like disease in patients with tick exposure^42,43^, was identified in all of the selected libraries (Figure 2). *Blattabacterium, Buchnera, Spiroplasma*, and *Candidatus Midichloria* were all present in at least 90% of samples, and *Candidatus Carsonella* was identified in 25% of samples (Figure 2). To our knowledge, none of these known endosymbionts have been commonly identified in *I. pacificus*, and *Blattabacterium, Buchnera*, and *Candidatus Carsonella* represent the first report of these genera in any tick^44,45^.

Sequencing individual ticks also provided sufficient resolution for co-occurrence analyses. We assessed whether presence of one microbial genus increases the statistical likelihood that another microbial genus will be present in the same tick host^46^. This revealed 14 pairs of bacterial genera detected together in a statistically significant number of samples (Figure S3). Of note, there was a strong positive association between the endosymbiont *Candidatus Carsonella* and *Chryseobacterium*, a genus that has been shown to be pathogenic to soft ticks but tolerated by hard ticks^47^. The remaining statistically significant co-occurrence relationships did not include any microbial genera known to be tick-associated, and we hypothesized that they may be environmental contaminants, such as soil bacteria. Therefore, outside of the *Candidatus–Chryseobacterium* case, we did not find clear evidence that any microbes associated with *I. pacificus* actively promote the colonization or growth of other microbes.

Our RNA-based sequencing approach also enabled identification of several previously unidentified RNA viruses in *I. pacificus* ticks. Ticks are known to carry a diversity of viruses, and the majority of known transmissible arboviruses have RNA genomes^15,48–52^. We sought evidence of these and any novel tick viruses in our metatranscriptomes. To do so, we first developed a bioinformatics strategy because standard tools for microbiome analysis (e.g., kraken2) did not detect any known tick viruses in our metatranscriptomic data^41^. This is a common phenomenon in RNA virus discovery, due to the fact that the diversity of RNA viruses are not well represented in reference databases. In keeping with other viral discovery efforts^53–55^, we searched for sequences containing an RNA-dependent RNA polymerase (RdRp) domain using HMMER^56^ (Figure 3a). Using this strategy, we detected a total of 13 new tick viruses in our *I. pacificus* field specimens and determined their prevalence across the dataset as well as their relative abundance within each sample (Figure 3b, Table 1). Underscoring the novelty of these viruses, all had less than 80% amino acid identity to their nearest relative in the NCBI non-redundant protein database (Figure 3c). While many of these viruses (10/13) could be defined as members of viral families previously identified in tick species, some had much less clear phylogenetic placements and no homology to known tick viruses. We named the viruses according to geographic features in the region in which the samples were collected.

### RNA viruses are important and stable constituents of tick microbiota

Our RNA-based approach to tick microbiome characterization led to the discovery of several novel viruses and viral families in *I. pacificus* field ticks. To follow up on these results, we next asked if the viruses we detected were likely to represent the full virome of the sampled ticks or only the most abundantly transcribed viruses. To do so we performed rarefaction analysis, which shows the number of new taxa discovered as a function of the number of samples sequenced. Our rarefaction curve appears to be approaching an asymptote, and the estimated true number of viruses in this population using the Chao index is 14.2 (Figure 3e). These results indicate that we have likely discovered the majority of viruses in this population with a median of 1.7 million non-host reads after DASH host-subtraction. This is equivalent to an overall sequencing depth of 106 million reads. We also observed that relatively few samples are needed to saturate viral discovery for a given population at this sequencing depth. Hence, we propose ~100 million RNA reads with DASH and rarefaction analysis as a standard for characterizing other arthropod viromes.

We also performed co-occurrence analysis with our newly characterized tick viromes. Not only did we identify a broad diversity of viruses, but we also found evidence of co-occurring viral infections within individual ticks. Ticks had a median of two viruses present with a maximum of six in one individual (Figure S4a). We found a statistically significant positive relationship between *Portuguese Ridge Virus* (Narnaviridae), *Wildcat Canyon Virus* (family unknown), and *Kasler Point Virus* (family unknown). Notably, all three of these viruses contained only an RdRp; no additional segments or genes were identified. Since Narnaviruses are single-gene ribonucleoprotein complexes lacking structural proteins or capsids, it is possible that the two viruses of unknown origin replicate and transmit in a similar manner. Our findings reinforce the model that ticks can harbor multiple viruses per individual, suggesting that RNA viruses are an important and stable part of the tick microbiome.

### Tissue and life-stage tropism of viruses suggests horizontal and vertical transmissibility

Arthropods are known to tolerate viral infection more easily than vertebrates, often maintaining infections for life with no apparent ill-effects^57^. Given that some of the viruses we identified are not only highly prevalent but also closely-related to viruses previously discovered in tick cell lines (Supplementary Note: *Rhabdoviridae*) ^26^, we next investigated how *I. pacificus* ticks may have acquired them. To explore whether any of the viruses we identified could represent viruses stably associated with *I. pacificus* ticks, we experimentally screened cDNA generated from pools of both wild and laboratory-reared *I. pacificus* larvae by polymerase chain reaction (PCR) using primer sets specific to all 13 of the identified viruses. Nine of the 13 viruses were present in either wild-collected or laboratory-reared larvae that have not yet consumed a bloodmeal, suggesting a likely scenario of vertical transmission (Figure 3d, Figure S4c).

We next screened laboratory-reared nymphs and adults to determine whether these viruses remain associated with ticks throughout life stages. Eight of the 9 viruses identified in larvae were also identified in either laboratory-reared nymphs or adults, suggesting these viruses persist through life stages and that they can be maintained in the population in the absence of their natural bloodmeal hosts (Figure 3d). We therefore investigated whether *I. pacificus* ticks could transmit their associated viruses to bloodmeal hosts via feeding. Microbes can be transmitted from tick midguts or salivary glands to their hosts via saliva secreted into the host bloodstream. We reasoned that viral presence in salivary glands in particular may correlate with feeding-based (horizontal) transmission (Figure S4c). We used our PCR assay to screen salivary gland and midgut cDNA libraries from additional field-collected *I. pacificus* ticks. The majority of the viruses identified (10/13) were detectable by PCR in tick salivary glands (Figure 3d). While this does not definitively demonstrate that these viruses are transmissible by feeding, we hypothesized that it could potentiate such a model.

To more directly probe whether the viruses we identified might be capable of infecting mammalian hosts, we tested for evidence of viral transmission to laboratory mice exposed to infected ticks. Because tick larvae must be ground up whole in order to extract RNA, the same tick cannot be tested at multiple time points. However, given that 6 of the viruses were identified in two pools of laboratory-reared larvae, we reasoned that additional pools from the same population were likely to contain the same viruses. We therefore fed two additional pools of laboratory-reared larvae on two mice to test whether any viruses present in the larval population were able to be transmitted to the mice. We performed PCR for the 13 viruses on blood samples collected from the mice before and during feeding, as well as the larvae after feeding (Figure S4d). Five viruses were detected in one or both mice throughout the experiment, in some cases even prior to feeding (Figure 3d, Figure S4e). This suggests that in addition to their association with *I. pacificus* ticks, these viruses may be commonly circulating in small mammals. In summary, our *in vivo* tick feeding experiments suggest that at least a subset of identified viruses are potential tick-borne viruses. Furthermore, our findings suggest that, contrary to our motivating assumptions, transmissibility is not strictly linked to stable salivary gland localization or vertical maintenance by *I. pacificus*.

### Identification of novel mRNA-like virus-like transcripts

In addition to the 13 viral RdRps, we identified 21 sequences with homology to an RdRp but with an open reading frame (ORF) structure inconsistent with known RNA viral genomes. Specifically, these RNA sequences encoded clusters of small ORFs with RdRp homology and large gaps (100s of bases) between their predicted ORFs (Figure 4a). Many also contained multiple overlapping ORFs and/or small ORFs in opposite orientations. These unusual sequences were highly prevalent (Figure 4a) and were independently assembled from multiple different ticks. Our analysis of these sequences suggested that they originated from many of the same families as the viral genomes (Figure 4b), but with distinct sequences that encoded ORFs that were smaller and more numerous than would be expected for that family.

**Figure 4:**
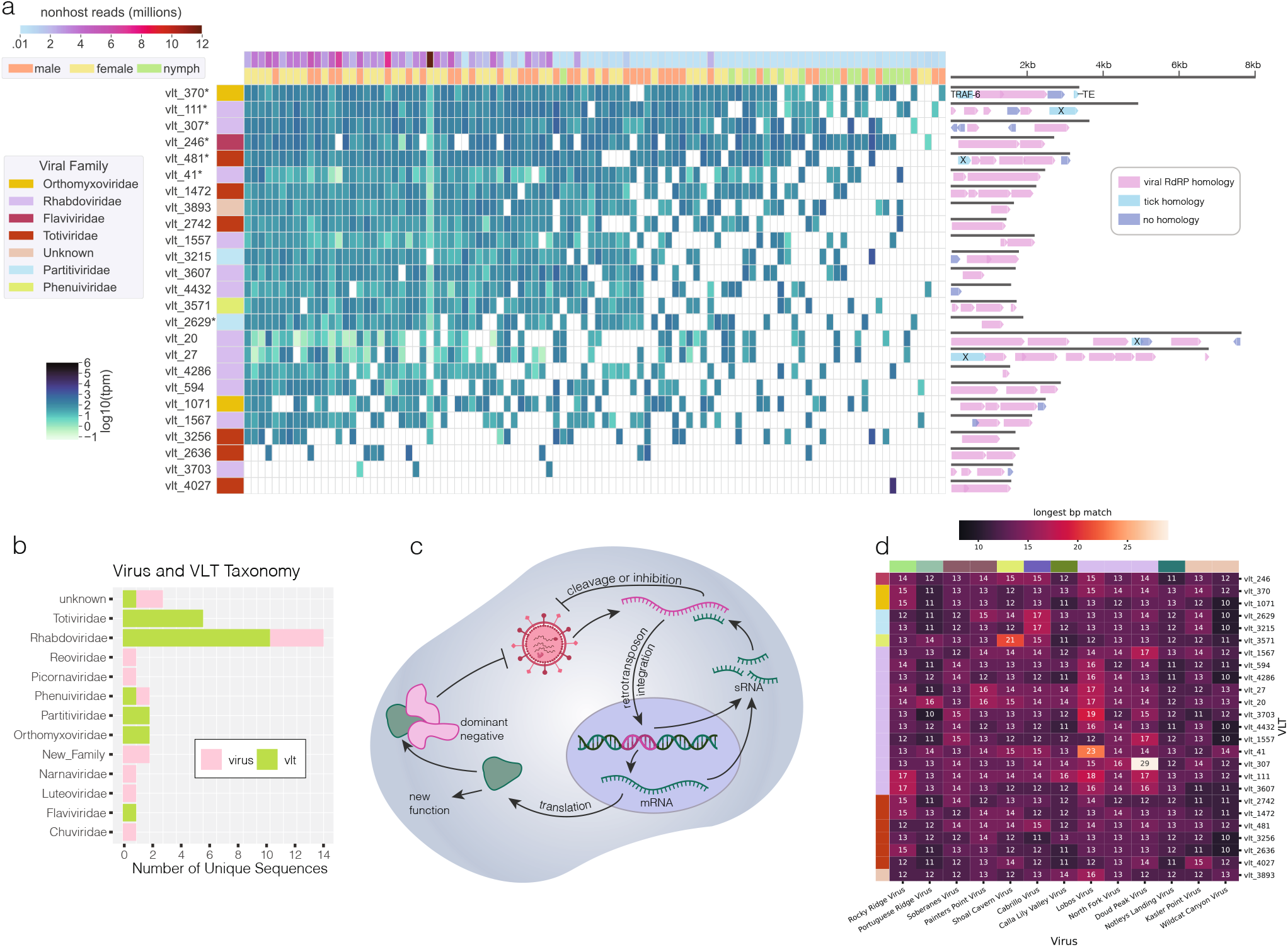
Virus-like transcripts detected across *Ixodes pacificus* population. a) Heatmap showing transcript per million value of virus-like transcripts (VLTs) across the dataset. Rows with an * were detected in DNA by PCR. Each VLT is colored by the viral family of its closest hit in blast and predicted open-reading frames (ORFs) are shown to the right. All ORFs in pink have homology to a viral RdRP, ORFs with homology to tick sequences are shown in blue and labeled by name, TRAF=TNF receptor-associated factor 6-like, TE=piggy-Bac transposable element-derived protein 4-like, X=protein of unknown function b) Viral family assignment of both exogenous viruses and virus-like sequences identified. c) Known functions of arthropod endogenous viral elements. d) Heatmap displaying length of longest perfectly matching sequence between each virus and each VLT. Rows and columns are colored by viral family.

We were intrigued by the observed irregular genomic organization of these 21 virus-like sequences; thus we next sought to better understand their possible origins and functions. We first conducted experiments to eliminate potential artifactual explanations for the irregular ORF structure. To test whether these sequences could be the result of a misassembly, we selected one of the longest and most highly prevalent sequences (vlt_111) for more in-depth evaluation. We applied RACE sequencing to examine the sequence in cDNA of a Garrapata tick. Our results confirmed the accuracy of the vlt_111 sequence assembly and indicated that vlt_111 is expressed as a 3’ poly-adenylated mRNA, ruling out the possibility that the non-canonical features of this RNA are due to misassembly of sequencing reads.

We also considered whether the irregular ORF structures we observed could be resolved with alternative codons, such as non-standard stop and start codons. We tested whether ORF prediction with any alternative genetic codes would result in an organization more consistent with that of a viral genome. Of the 25 known genetic codes tested, none substantially changed the ORF structure of any of the sequences (Figure S7d), indicating that alternative genetic code alone cannot account for the observed genomic structure. Having eliminated possible artifactual explanations for how these sequences could originate from exogenous viral genomes, we termed these sequences of unknown origin and function “virus-like transcripts” (VLTs).

### VLTs likely serve a non-canonical function in I. pacificus ticks

Arthropod genomes are known to contain numerous endogenous viral elements (EVEs), which result from horizontal integration of RNA viral sequences into tick genomes over the course of evolution^28^. The observed high prevalence of these VLTs lead us to hypothesize that these mysterious sequences could have likewise arisen from such genomic integration events. In the absence of a published genome assembly for *I. pacificus*, we could not check for corresponding sequences in a reference genome. Therefore, we developed PCR primers to screen *I. pacificus* DNA extracts isolated from the same wild-collected ticks used for metatranscriptomics for VLTs. We confirmed the presence of vlt_111 in wild-collected tick DNA (Figure S7b) as well as laboratory-reared tick DNA. To ensure this pattern was not specific to this particular VLT, we checked an additional 5 VLTs, all of which were present in lab-reared *I. pacificus* DNA (Figure S7c), suggesting a genomic origin for these VLTs.

To confirm that the presence of DNA forms of transcripts is specific to the VLTs and not a general phenomenon, we additionally screened all of the presumed exogenous viral genomes for presence in genomic DNA. Only one of the viral genomes was detected in DNA. A faint band corresponding to *Rocky Ridge Virus* was amplified from genomic DNA. This virus, a mivirus with a circular genome, could represent an intermediate between these two categories that is an exogenous RNA virus with a single or small number of recent genomic integrations into the I. pacificus genome. Alternatively, this could be caused by production of DNA forms of the viral genome by endogenous retrotranscriptases (either prior to genomic integration or in the absence of integration) as in Salvati et al^58^. Given that its expression pattern mirrored that of the other exogenous viruses and its genome is both complete and contains the expected ORFs, we continued to classify this sequence as an exogenous virus and not a VLT.

To explore the possibility that our VLTs could have EVE-like functions, we next considered canonical pathways by which EVEs contribute to arthropod immunity (Figure 4c). EVEs are most commonly known to function as non-coding RNAs. Much more rarely, they are expressed and translated as proteins that can act as dominant negative viral inhibitors or serve a new function^28,29^. As the fragmented ORF structure of our VLTs is inconsistent with expression of full-length proteins, we focused on non-coding RNA functions. Typically, arthropod EVEs play antiviral roles by serving as a template for piwi RNAs (piRNAs), 24–31 nucleotide (nt) RNAs that target an exogenous viral RNA genome for degradation by binding to a complementary sequence within it.

To test this model, we examined *in silico* whether our VLTs could give rise to small RNAs capable of binding the exogenous viral genomes in our dataset through a matching sequence at least 24 nt long. Only one VLT contained a sequence of at least this length (vlt_307) matching one of our viral genome assemblies. Two others contained stretches longer than 20 nucleotides (Figure 4e). The remaining 18 VLT sequences do not contain perfect matches longer than 17 nucleotides to any of the exogenous viruses. Of the three VLT-virus combinations with perfect matches of at least 20 nucleotides, none originated from the same tick sample (Figure S8). In total, we did not uncover definitive evidence supporting inhibition of viral replication through a canonical piRNA pathway. Our results strongly point to either a non-canonical immunity mechanism or a different functional role entirely for the VLTs we identified in *I. pacificus*.

## Discussion

In this study, we show the power of combining experimental enrichment of microbial sequences with single-tick metatranscriptomics for identification of bacteria and viruses in field *I. pacificus* tick communities. The ability to deeply sequence the non-host fraction allowed us to identify several genera of bacteria in *I. pacificus* previously unidentified in any tick species. Further investigation is warranted into the consequences and mechanisms underlying these symbiotic tick–bacteria partnerships.

Our approach also enabled us to uncover many novel viruses, which we further investigated in a series of laboratory experiments. We found that several of these viruses were not only highly prevalent but were also present in tick salivary glands. This has important implications for public health as bloodmeal hosts (including humans) are very likely to be exposed to viruses present in the salivary glands during feeding. The detection of these viruses in mice further suggests they are circulating between ticks and mammals. Currently, tick-borne disease surveillance in California is focused on Lyme disease and a small number of other bacterial pathogens, but these results indicate patients should also be screened for viruses^1^.

In addition to providing more comprehensive and quantitative insights into the *I. pacificus* microbiome, one of the most exciting and unexpected themes that emerged from our work relates to how viruses likely play fundamental and important roles in tick biology. We found evidence that several viruses persisted in ticks across multiple life stages, including juvenile naïve larvae, as well as across wild and laboratory-reared populations, suggesting they are stable constituents of the tick microbiome. Certain viruses in *I. pacificus* may be able to establish and maintain independent niches within their tick hosts. These findings lay the groundwork for future work aimed at understanding tick–virus dynamics and how such relationships may play fundamental roles in tick physiology.

Further underscoring the critical importance of tick–virus interactions for *I. pacificus* biology was our discovery that numerous VLTs may be integrated in the tick genome as EVEs. EVEs may be an underexplored feature of tick genomes that can be identified in future studies through tick genome studies or an RNA sequencing-based approach such as ours that enriches for low abundance transcripts. Evidence for horizontally acquired EVEs in *I. pacificus* raises the possibility that ticks have hijacked viral sequences and functions for some adaptive benefit provided to the tick. Closer experimental evaluation of our VLTs pointed to a non-canonical mechanism that is distinct from known antiviral pathways. The VLTs we found may be unique to or enriched in ticks, providing a useful handle for interrogating a new mechanistic class of EVEs that may have important contributions to tick immunity and biology.

The integration of RNA viral genomes into the tick genome as EVEs also provides a unique historical footprint for viruses that may have infected that tick host in the near or distant past. Interestingly, the VLTs identified in *I. pacificus* appear to derive from several viral families from which no exogenous viral genomes were found in this study, including the recently-discovered segmented flaviviruses that cause febrile illness^59–61^. Future field studies that expand on our *I. pacificus* virome analyses will help determine whether VLTs stem from ancient tick–virus interactions or contemporaneous interactions that were not captured in this study due to low abundance, our limited sample size, and our focus on two collection locations. Altogether, our results highlight the need for more studies such as this in order to capture the full range of tick–associated microbes that could represent critical components of tick physiology or poorly understood pathogenic threats to human health. Our work provides an improved experimental and computational framework with increased sensitivity for low-abundance bacterial and viral taxa present in this increasingly important class of arthropod disease vectors.

## Methods

### Tick Source

Ticks for the initial metagenomic sequencing were collected by dragging from Garrapata State Park (n=83) and China Camp State Park (n=17). Ticks for followup laboratory experiments were collected exclusively from Garrapata State Park. Adult ticks were separated by sex, surface sterilized in 1% bleach and frozen individually.

Laboratory-reared I. pacificus were received from the tick lab at the Center for Disease Control tick lab (Atlanta, GA) and provided through BEI Resources (a service funded by the National Institute of Allergy and Infectious Diseases and managed by ATCC). Ticks were maintained in glass jars with a relative humidity of 95% (saturated solution of potassium nitrate) in a sealed incubator at 22°C with a light cycle of 16h/8h (light/dark).

### RNA Extraction/Library Prep

Total RNA was extracted from the wild-caught *I. pacificus* adult and nymph ticks in 2 separate batches. On ice, individual ticks were transferred to separate wells of a 96 well deepwell plate that was pre-loaded with a single 5mm steel ball bearing (OMNI International, GA, USA) and 400uL of 1X DNA/RNA shield (Zymo Research Corp., Irvine CA, USA) in each well. The plates were sealed and subjected to bead bashing (3 x 3 min, with 1 min rest on ice in between each round of bashing) on a TissueLyser II beadmill (Qiagen, Valencia, CA, USA), then clarified by centrifugation at 2000 rpm at 4°C for 5 min in a refrigerated tabletop centrifuge (Beckman Coulter, Indianapolis IN, USA) to remove large debris. 350uL of the supernatant was transferred to a fresh 96 deepwell plate and re-centrifuged under the same conditions to further clarify the homogenate. 90uL of the resulting supernatant was used as input for total RNA extractions; 110uL of supernatant was transferred to a separate plate and archived at −80°C for potential follow-up analyses.

For both the adult tich and nymph tick homogenate preps, automated RNA extraction was performed in 96 well format (Bravo automated liquid handler, Agilent Technologies, Santa Clara, CA, USA) using a modified version of the Quick DNA/RNA pathogen magbead 96 extraction kit (Zymo Research Corp., Irvine, CA, USA) to automate total nucleic acid extraction and DNase treatment. RNA extracted from 90 uL of tick homogenates was eluted in a final volume of 25uL into 96 well PCR plates. An aliquot of 3uL was used for quantitative and qualitative analysis of the total RNA for each sample via Qubit fluorometer assay (Thermo Fisher Scientific, Waltham MA, USA) and Agilent Bioanalyzer Pico 6000 total Eukaryotic RNA electrophoresis (Agilent Technologies, Santa Clara, CA, USA). A separate 5uL aliquot was used as input for RNAseq library prep, and 2 x 7uL aliquots were stamped into 2 separate daughter plates that were immediately frozen and archived at −80°C for potential follow-up studies.

RNAseq libraries preparation of the 5uL aliquots of adult tick and nymph tick total RNA preps was also performed in 96 well format on an automated liquid handler (Bravo automated liquid handler, Agilent Technologies, Santa Clara, CA, USA). Briefly, the NEBNext Ultra II Directional RNAseq library preparation kit (New England Biolabs, Ipswich, MA, USA) was applied with the following modifications incorporated into the manufacturer’s standard protocol: a 25pg aliquot of External RNA Controls Consortium RNA spike-in mix (“ERCC”, Thermo-Fisher, Waltham, MA, USA) was added to each sample prior to RNA fragmentation; the input RNA mixture was fragmented for 8 min at 94°C prior to reverse transcription; and a total of 12 cycles of PCR for amplification of resulting individual libraries. SPRIselect (Beckman Coulter, Indianapolis IN USA) beads were used to size-select libraries with an average total length between 450–550 bp. Library size distributions were verified by Agilent Bioanalyzer High Sensitivity DNA electrophoresis (Agilent Technologies, Santa Clara, CA, USA) and quantified by Qubit fluorometer (Thermo Fisher Scientific, Waltham MA, USA). Paired-end 2 x 150bp sequencing runs were performed on equivolume pools of individual sequencing libraries of the adult and nymph ticks, respectively, on the Illumina MiSeq sequencing platform (Illumina, San Diego, CA, USA).

The yield of reads/uL acquired from the small scale MiSeq run of the equivolume pools of individual libraries were used to generate approximately equimolar pools of the individual adult tick and nymph tick libraries. The pooled libraries were then depleted of highly abundant sequences^40,62^, using a previously described pool of tick gRNAs^63^ complexed with in-house prep of purified recombinant Cas9 protein. Resulting DASH’d libraries were qualitatively and quantitatively analyzed by Agilent Bioanalyzer High Sensitivity DNA electrophoresis (Agilent Technologies, Santa Clara, CA, USA) and Qubit fluorometer (Thermo Fisher Scientific, Waltham MA, USA). While the DASH’d adult tick libraries provided sufficient material for large scale metatranscriptomic sequencing, insufficient material remained in the tick nymph libraries that were DASH’d. Thus for large scale metatranscriptomic sequencing, the pool of DASH’d adult tick libraries was combined with a pool of un-DASH’d nymph tick libraries. This pooled prep was subjected to PE 2x150bp format on the NextSeq2000 Illumina sequencing platform (Illumina, San Diego, CA, USA).

### Host Subtraction/Pre-Processing

Fastq reads from the run were pre-processed using the CZID pipeline. Libraries underwent quality filtering and adaptor trimming. Host reads were then removed by mapping to closely related genomes. Reads from each library were then assembled into contigs using SPADEs^64^ within the CZID pipeline^65^. Both the nonhost reads and resulting contigs were used for downstream analysis.

### Bacterial Classification

Host-subtracted reads were classified using Kraken2 (version 2.1.1)^41^. The full kraken2 database was used for classification. Only libraries with at least 1000 classified reads were considered for analysis. Reads per taxon were converted to reads per million (rpm) using library size. To reduce false positives, the rpm value for each taxon was required to be at least 100 times the rpm in any of the control libraries (water and Hela cells) to be considered a positive. Additionally, at least 100 unique minimizers were required for each taxa. Taxa within each library not meeting these thresholds were excluded from analysis. Fewer genera were detected in smaller libraries, including the nymphal ticks from China Camp SP (Figure S1a), and samples clustered primarily by library size. We therefore focused subsequent analysis on libraries of at least one million non-host reads.

### Virus Identification

We focused our analysis on RNA viruses, as these tend to dominate arthropod viromes^32,66,67^. Contigs were filtered to those of at least 1500 base pairs. Open reading frames were predicted for these contigs using prodigal^68^, and the resulting proteins were searched using HMMscan from HMMER3 (version 3.3.2)^56^ against a collection of HMMR profiles of viral RdRPs. The following RdRP HMMs were downloaded from the pfam database^69^ on March 4, 2021; RdRP_1, RdRP_2, RdRP_3, RdRP_4, RdRP_5, Viral, RdRp_C, Mitovir_RNA_pol. Mononeg_RNA_pol, Birna_RdRp, and Bunya_RdRp. Additionally, custom HMMs were constructed from RdRP sequences for narnaviridae and orthomyxoviridae (sequences and combined HMM available in supplement).

Any sequences with a putative RdRP hit from HMMER were then queried against the full NCBI nonredundant protein database (as of January 24, 2021) using diamond blastp version 0.9.24 to identify their closest hit^70^. For proteins with multiple hits, the hit with the highest bitscore was reported. Sequences covering less than 30% of their closest blast hit were initially classified as putative virus-like transcripts, and remaining sequences were classified as exogenous viral sequences. The open reading frame (ORF) structure of all sequences was then manually inspected to confirm this classification. Sequences containing significant gaps between ORFs, or multiple reading frames with RdRP homology (where one was expected) were further classified as virus-like transcripts (VLTs).

### Determination of Prevalence and Abundance

Assembled contigs were clustered using cd-hit-est (CDHIT version 4.8.1) at a threshold of 85% nucleotide identity^71^. Circular chuvirus genomes were rotated to a common start position using a custom python script to ensure accurate clustering. 85% identify was chosen as a cutoff to minimize multi-mapping reads between closely related sequences. However, some clusters contain significant sequence diversity and could potentially be considered to contain multiple species. The representative sequences from this clustering were used for all downstream analysis.

Reads from each library were mapped back to the collection of cluster representatives using bowtie2 version 2.4.1^72^. Reads aligning to each contig were counted using samtools idxstats version 1.9^73^. Aligned reads, contig length, and library size were used to calculate rpm and transcripts per million (tpm) values for each library. To consider a contig “present” in a given library, the rpm value was required to be greater than 10 times the value in any of the control libraries. This filter was designed to remove potential false positives caused by cross-contamination of high-titer species.

### Identification of Additional Genomic Segments

To identify additional segments of multipartite viral genomes, we searched for contigs that were strongly co-occurring with the RdRp containing contigs. Presence/absence of each contig cluster for each library was determined by reads mapped to each contig using the filters described above. Presence was coded as a 1 and absence as a 0 and the Jaccard distance was calculated for all pairs of contigs. We considered any sequence with Jaccard distance < 0.4 as a putative genomic segment and further considered homology of the sequence and whether additional segments are expected for the viral family in our determination of whether these sequences represent segments from the same genome (Figure S6).

### Rarefaction Analysis

The viral genomes in each sample were determined by the presence of a contig of at least 1000 base pairs (bp) that clustered with one of the 13 representative genomes identified. The presence of a contig rather than read mapping was used to simulate viral discovery in each sample, under the assumption that new viruses discovered may be too divergent to detect by read mapping. The samples were ordered by the number of new genomes seen (not seen in any of the previous samples). The number of new genomes was counted for the addition of each sample. This was repeated for 50 iterations and the median number of new samples at each step was determined. The Chao index was calculated using the R library fossil version 0.40^74,75^ .

### Co-Occurrence of Taxa

To determine whether any pairings of taxa (either bacterial or viral) occur more or less frequently than expected given their prevalence we utilized the recently developed metric ***α***^46^. The presence of each taxon was considered at the genus level for bacteria and at the species level for viruses. The distance ***α*** and associated p-value were determined for all pairs of taxa using the CooccurrenceAffinity R package (version 1.0)^46^. Pairs were filtered to those with a p-value ≤ 0.005. These relationships were visualized as a network with edges corresponding to ***α*** and nodes corresponding to taxa using the R package igraph (version 1.3.0)^76^. Node size was scaled according to taxon prevalence in the dataset.

### Phylogenetic Trees

Trees were constructed using the RdRP protein sequence of each virus identified. The top 100 closest blast hits were downloaded for each virus and clustered at 85% nucleotide identity. Sequences were aligned using MAFFT version 7.475^77^ and maximum likelihood trees were constructed using iqtree 2.0.3^78^. Trees were visualized in iTOL version 5^79^.

### Virus-like Sequence PCR

To verify that virus-like transcripts were present in tick genomes and expressed in wild and lab-reared ticks, we extracted tick RNA and genomic DNA and confirmed the presence of virus-like transcripts with PCR. Adult male *I. pacificus* ticks (n=3) were pooled and homogenized by beating for 2 increments of 30 seconds at 4000 bpm in a bench homogenizer (Bead Bug, Benchmark Scientific) with 1.4mm zirconium oxide ceramic beads (Fisher Scientific) in ice-cold TRIzol reagent (Thermo Fisher Scientific). RNA extraction was performed using a Zymo Research Direct-zol RNA Microprep kit (Zymo Research), and RNA was converted to cDNA using Primescript RT reagent kit (Takara Bio) in 10 uL reactions following manufacturer protocols. Genomic DNA was extracted from adult male *I. pacificus* ticks (n=3, separate individuals from RNA), which were pooled, flash frozen, and ground to powder. A DNeasy Blood & Tissue Kit (Qiagen) was used to extract genomic DNA following manufacturer protocols.

PCR experiments amplifying regions of VLTs from tick cDNA and genomic DNA were run using Platinum Superfi II Green PCR master mix (Thermo Fisher Scientific). We loaded 5uL of product onto 0.7% agarose gels and ran them at 160V for 2 hours before imaging. Primer sequences can be found in Table 2.

### Virus multiplex PCR

Viral sequences were analyzed with Snapgene (Dotmatics) and PrimerPlex software (Premier Biosoft) to design 4 sets of multiplex primers that amplify 100–550bp regions. Platinum™ SuperFi II Green PCR Master Mix (Thermo Scientific) was used for all PCR reactions. Mixed cDNA from the original sequenced field-collected ticks was used as a positive control. No-template (water only) reactions were also included as negative controls. Primer pair sequences are listed in Table 2. PCR reactions were analyzed by electrophoresis using a 2% agarose gel containing GelRed (Biotium) and visualized using an Azure c400 imager (Azure Biosystems). At least one band corresponding to each virus was cut out of the agarose gel, purified using the QIAquick Gel extraction kit (Qiagen), and sanger sequenced (GeneWiz) to confirm correct PCR amplification of the intended virus.

### Mouse Feeding Experiment

Animal experiments were conducted in accordance with the approval of the Institutional Animal Care and Use Committee (IACUC) at UCSF. *I. pacificus* larvae (CDC/BEI) were fed on young female C3H/HeJ mice acquired from Jackson Laboratories. Mice were anesthetized with ketamine/xylazine before placing ~50 larvae. Ticks allowed to feed to repletion and collected from mouse cages. Whole blood was collected from mice before (pre), during (Day 2), and at the end (Day 4) of larvae feeding to access viral transmission by PCR (see above).

### Tick Dissections/Extractions

For tissue-tropism determination, wild-collected ticks (n=20) from Garrapata State Park were dissected using a micro scalpel cleaned with 70% isopropanol and a sterile needle. Ticks were dissected in batches of 3–5. The scalpel was cleaned and the needle was replaced between batches. The tick cuticle was excised and the midgut and salivary glands were removed using tweezers cleaned with 70% isopropanol. Tissues were pooled and rinsed in droplets of PBS then transferred by pipette into 300uL of Trizol. Males and females were processed separately and each pool of tissues contained material from 3–10 individuals.

Whole adult ticks were added to 300uL of Trizol in pools of 4–5 individuals, grouped by species and sex. Nymphal ticks were added to Trizol in a pool of 3, and larval ticks were flash frozen and added to Trizol in pools of 10–15. All ticks and tick tissues were homogenized by bead-beating with ceramic beads in increments of 30 seconds. Samples were placed on ice between cycles and cycles continued until tissue was visually homogenized.

RNA extraction was performed using Directzol RNA Extraction kits, with on-column DNAse1 treatment. RNA was converted to single stranded cDNA using Quantabio cDNA mastermix in 10uL reactions.

## Acknowledgements

We would like to thank Anne Sapiro, Gytis Dudas, Kishen Patel, and Raul Andino for their helpful conversations and feedback on this project. We are also grateful to Greg Huber, Olga Botvinnik, Norma Neff, and CZB for their strategic and technical input. We also thank members of both the Chou lab and the lab of Andrea Swei (SFSU) for assistance collecting field ticks. We greatly appreciate all members of the Chou and Pollard labs for their input and feedback on experiments, analyses, and manuscript preparation.

## Supplementary Note

### Rhabdoviridae

*Rhabdoviridae* was the most common family identified. We detected three new members of this family, each containing either four or five open reading frames. *Lobos Virus* and *Doud Peak Virus* both group phylogenetically near other rhabdoviruses identified in ticks (Figure S5), including two viruses identified in Australian ticks that were not previously assigned to a clade^50^. These two viruses rarely occur in the same samples. Although this co-exclusion is not significant, it could indicate that these related viruses compete to occupy a similar niche in their host. Both of these viruses were confirmed by PCR in both salivary glands and midguts, as well as in wild-collected larvae and laboratory-reared adult females.

*North Fork Virus* was identified in 15% of samples. Its phylogenetic location is on a branch containing other viruses identified in ticks (Figure S5). Its closest relative is an endogenous virus that was discovered in the *I. scapularis* cell line IDE8, which causes no apparent cytopathic effect, raising the possibility that *North Fork Virus* may be a related endogenous virus of *I. pacificus*^59^. In keeping with this hypothesis, *North Fork Virus* was identified in a pool of laboratory-reared *I. pacificus* larvae by PCR, suggesting that it is able to be transmitted directly to offspring (who have not yet been exposed to an animal host).

### Bunyavirales

One phlebovirus was identified, which was named *Shoal Cavern Virus*. In addition to RdRP, a putative nucleoprotein segment was identified by co-occurrence. Phleboviruses are members of the *Phenuviridae* family and include tick-borne viruses known to cause human disease such as *Severe Fever with Thrombocythemia Virus* and *Heartland Virus*. *Shoal Cavern Virus* (*Phenuiviridae*) was not only highly prevalent (40%) but was also present in astonishingly high levels in some libraries, accounting for 38% of the total nonhost library in a single sample. This mirrors previous finding of a phlebovirus discovered in the hard tick *Dermacentor occidentalis* in California, adding further evidence that such viruses may be common tick endosymbionts^15^. *Shoal Cavern Virus* was identified in both salivary gland and midgut tissue, and it may therefore be transmissible to bloodmeal hosts, although its presence in wild larvae suggest it may also be vertically transmitted.

Two additional viruses were discovered that appear to group with other members of *Bunyavirales* but outside of known families. Phylogenetic analysis indicates they are most closely related to viruses of the *Hantaviridae* family (Figure S5). *Soberanes Virus* and *Painter’s Point Virus* each contain a single large, 9-kilobase (kb) open reading frame. A glycoprotein-encoding genomic segment was identified for each of these viruses, as well as one (*Painter’s Point Virus*) and two (*Soberanes Virus*) genomic segments of unknown function. Soberanes virus has no evidence of vertical transmission, as it was identified only in wild collected adult ticks (Figure 3d). It was furthermore detected in mice both before and during tick feeding, as well as in larvae fed on those mice, suggesting it could be transmitted horizontally between small mammals and ticks. *Painter’s Point Virus* was identified in wild samples as well as laboratory adults, nymphs, and larvae, suggesting the ability to be vertically transmitted. It was also detected in laboratory mice before and during tick feeding.

### Chuviridae

*Rocky Ridge Virus* belongs to *Chuviridae* and is composed of a 10.8 kb circular genome with three open reading frames. Since the discovery of this family, a number of chuviruses have been identified across continents and several tick families and have also frequently been found as endogenous viral elements in mosquito genomes^80^. The chuvirus detected in our study groups most closely with *Suffolk Virus*, a virus identified in *I. scapularis*. *Rocky Ridge Virus* was both highly prevalent (39%) and the most abundant virus of any in the dataset; nine samples contained more than 100,000 reads per million mapping to it. *Rocky Ridge Virus* was identified in nearly all tick samples screened by PCR, and its presence in salivary glands, and mice during feeding is evidence it may be transmissible to hosts, while its presence in larvae suggests it can also be vertically transmitted.

### Narnaviridae

*Portuguese Ridge Virus* is most closely related to viruses from the *Narnaviridae* family (Figure S5). Narnaviruses are unique in that they lack any structural proteins or capsids, existing instead as ribonucleoprotein complexes which are transmitted directly cell-to-cell either vertically or sexually. While first identified as viruses of fungi, they have since been identified in a variety of arthropods, sometimes with additional segments or ambigrammatic open reading frames^81^. *Portuguese Ridge Virus* was the only virus in the dataset to exhibit clear tissue tropism, identified only in midguts. It was additionally detected in mice both prior to and during tick feeding, which could explain its presence in the tick midgut (as a virus that can be acquired by but not transmitted by ticks).

### Reoviridae

*Calla Lily Valley Virus* is a Coltivirus in the family *Reoviridae* (Figure S5). Reoviruses have double stranded RNA genomes composed of up to 12 segments, and they infect a broad range of hosts including fungi, invertebrates, vertebrates, and plants. We identified an additional 8 segments by co-occurrence, four of which have homology to other *Reoviridae* proteins and four of which have no homology to known proteins (Figure 3a, Figure S6). Several members of this family have recently been identified in ticks and they are one of the most common families of endogenous tick viruses^26,82,83^ .

### Solemoviridae

*Notley’s Landing Virus* groups closely with other viruses identified in ticks in the now defunct *Luteoviridae* family (Figure S5). Interestingly, despite the high read coverage of this genome, it is smaller than expected, with viruses of this family typically being 5–6 kb in length with six open reading frames. This could indicate that Notley’s Landing Virus represents a new related family with a segmented genome, however no additional segments were identified by co-occurrence. *Notley’s Landing Virus* was not detected by PCR in either midguts or salivary glands but it was detected in laboratory-reared larvae, indicating it could be vertically transmitted.

### Picornaviridae

*Cabrillo Virus* is a member of *Picornaviridae*, a family of monopartite ssRNA viruses of genome size 7–9 kb encoding a single polyprotein. It is most closely related to *Falcovirus A1*, a virus identified in the common kestrel^84^. It is possible that *Cabrillo Virus* may be an avian-infecting virus as *I. pacificus* are known to feed on birds^85^. It was identified in nearly all tick samples tested by PCR, as well as a mouse during tick feeding, indicating both horizontal and vertical transmission.

### Unknown Family

Two additional genomes containing an RdRp were identified which do not have any homology to known sequences. Due to their small size, *Wildcat Canyon Virus* and *Kasler Point Virus* likely represent either segmented viruses or partial genomes. Although they could not be assigned to any known viral clade, they have 80% amino acid identity to each other and therefore likely belong to the same as yet unknown family. While *Kasler Point Virus* was identified by PCR in both wild-collected and laboratory ticks (including larvae) and mouse samples, *Wildcat Canyon Virus* was only identified in wild ticks, indicating potentially different modes of transmission. Interestingly, despite its presence in mice several contigs that co-occurred with *Kasler Point Virus* had homology to plant sequences, suggesting that it could be a plant virus that was sequenced on the outside of the tick.

## Supplementary Materials

Sequencing data is in submission process to the NCBI Sequence Read Archives and will be registered under project <u>PRJNA870442</u>

Intermediate files, including assembled contigs and read coverage can be found on figshare at https://figshare.com/projects/I_pacifics_mNGS/144081

Viral genomes are in submission process to NCBI genbank, they can currently be accessed under the above figshare using DOI 10.6084/m9.figshare.20497227

Analysis notebooks and scripts can be found on github at https://github.com/callamartyn/ipac_virus

CZID pipeline results, including intermediate host subtracted reads, assemblies, and other intermediate files can be found at https://czid.org/pub/uuFkacq3hT (Garrapata adults), https://czid.org/pub/EEJfJPNhYP (Garrapata adults, resequencing of low-coverage samples), and https://czid.org/pub/ss3AnxpDbU (China Camp nymphs)

---

**Figure S1:**
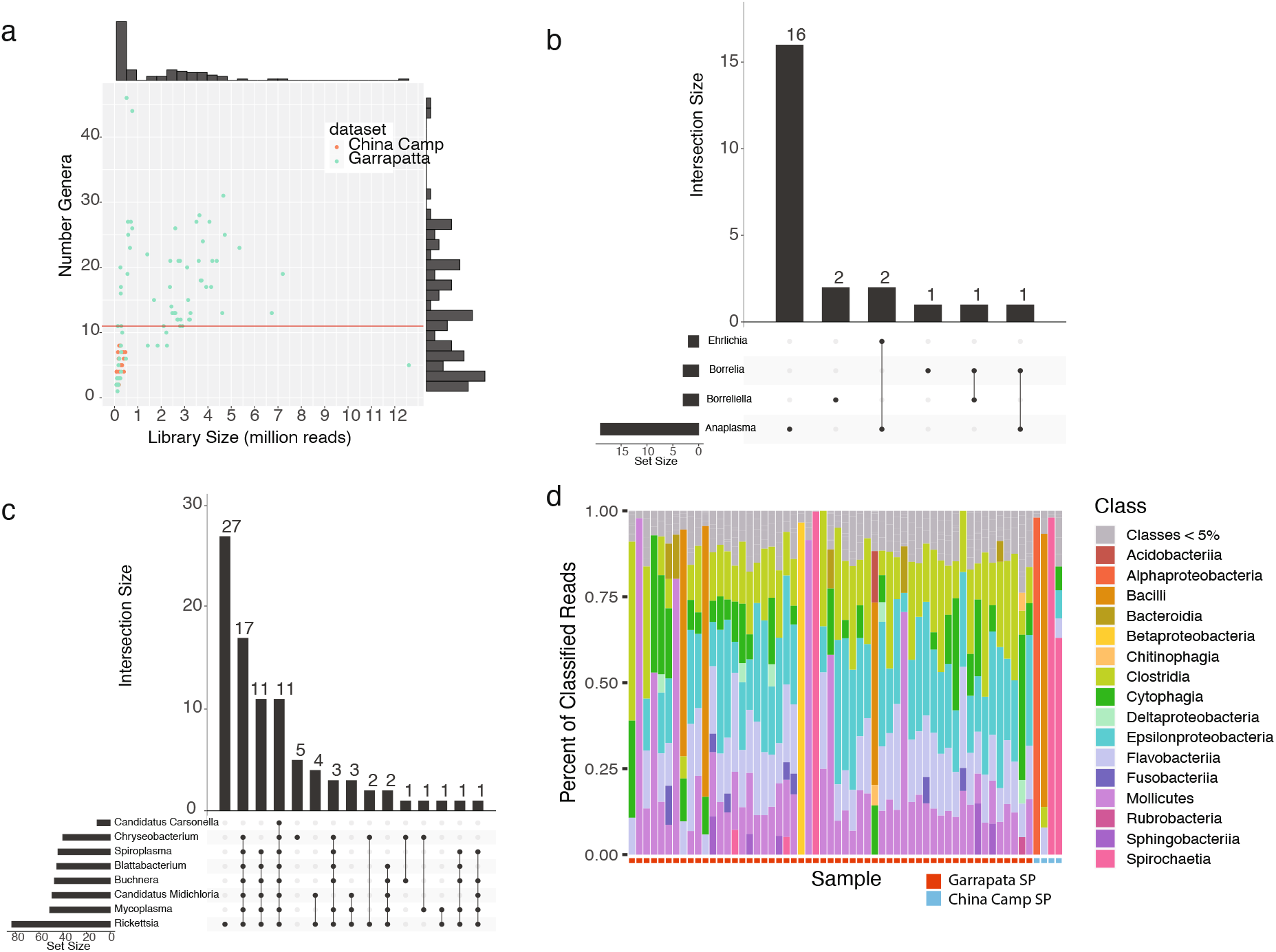
Pathogens and commensals of *I. pacificus*. a) Scatterplot displaying number of nonhost reads and number bacterial genera detected per sample. Horizontal line shows median number of genera across all samples. Histograms on each axisrepresentdistribution of axis values. b)Upset plot displaying number of coinfections of known bacterial pathogens. Numbers represent number of samples with the included set of genera. c) Upset plot displaying number of coinfections of newly identified endosymbionts. d) Strip plot displaying the propotion of reads assigned to each bacterial class by sample.

**Figure S2:**
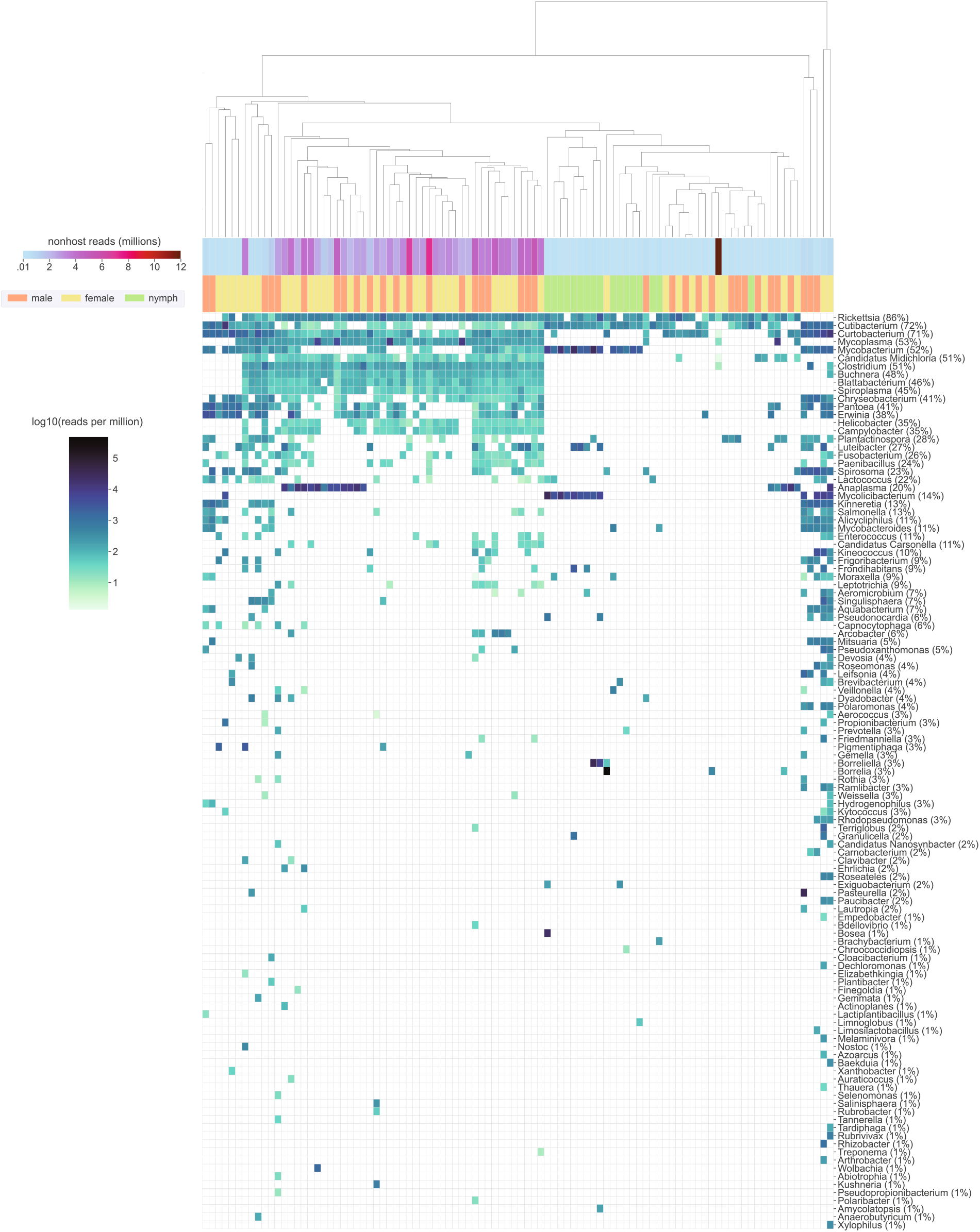
Bacterial genera of *I. pacificus*. Heatmap showing reads per million (rpm) of bacterial genera as classified by kraken2. Rows are ordered in decreasing prevalence (shown next to genus name) and columns are heirarchically clustered by euclidean distance.

**Figure S3:**
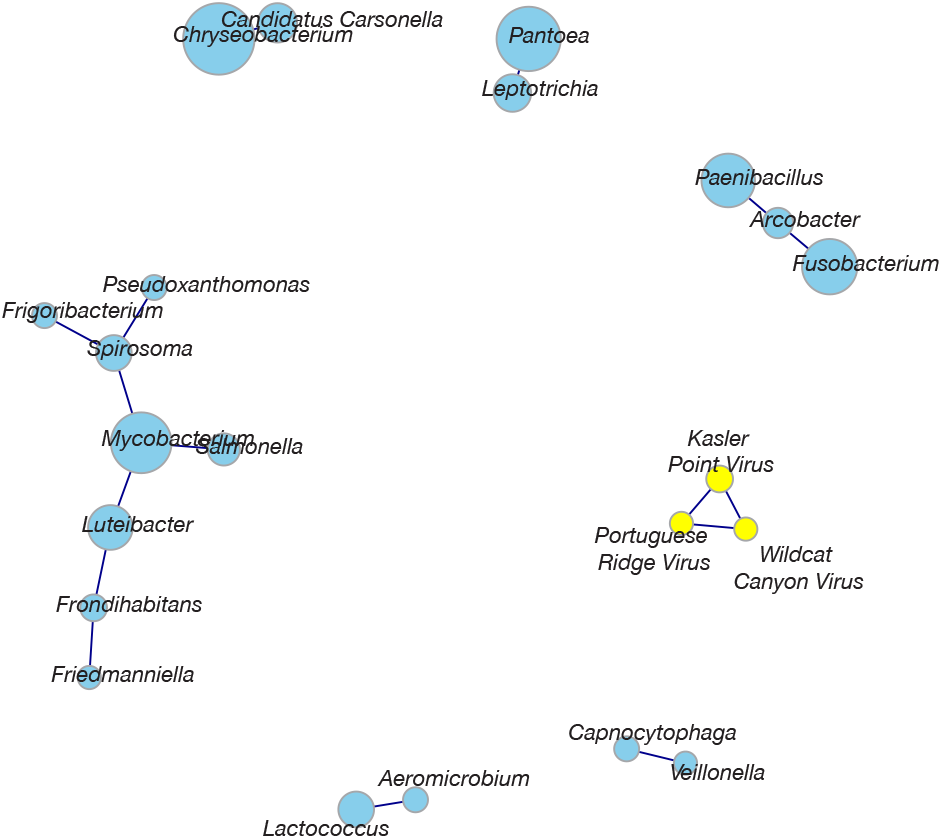
Co-occurence of tick microbes. Co-occurrence of bacterial and viral taxa network representation of significant co-occurring relationships amongst all identified viruses and bacterial genera. Sizes of nodes are scaled to the prevalence in the dataset. All edges represent a positive co-occurence of alpha value greater than or equal to 5 with a p-value less

**Figure S4:**
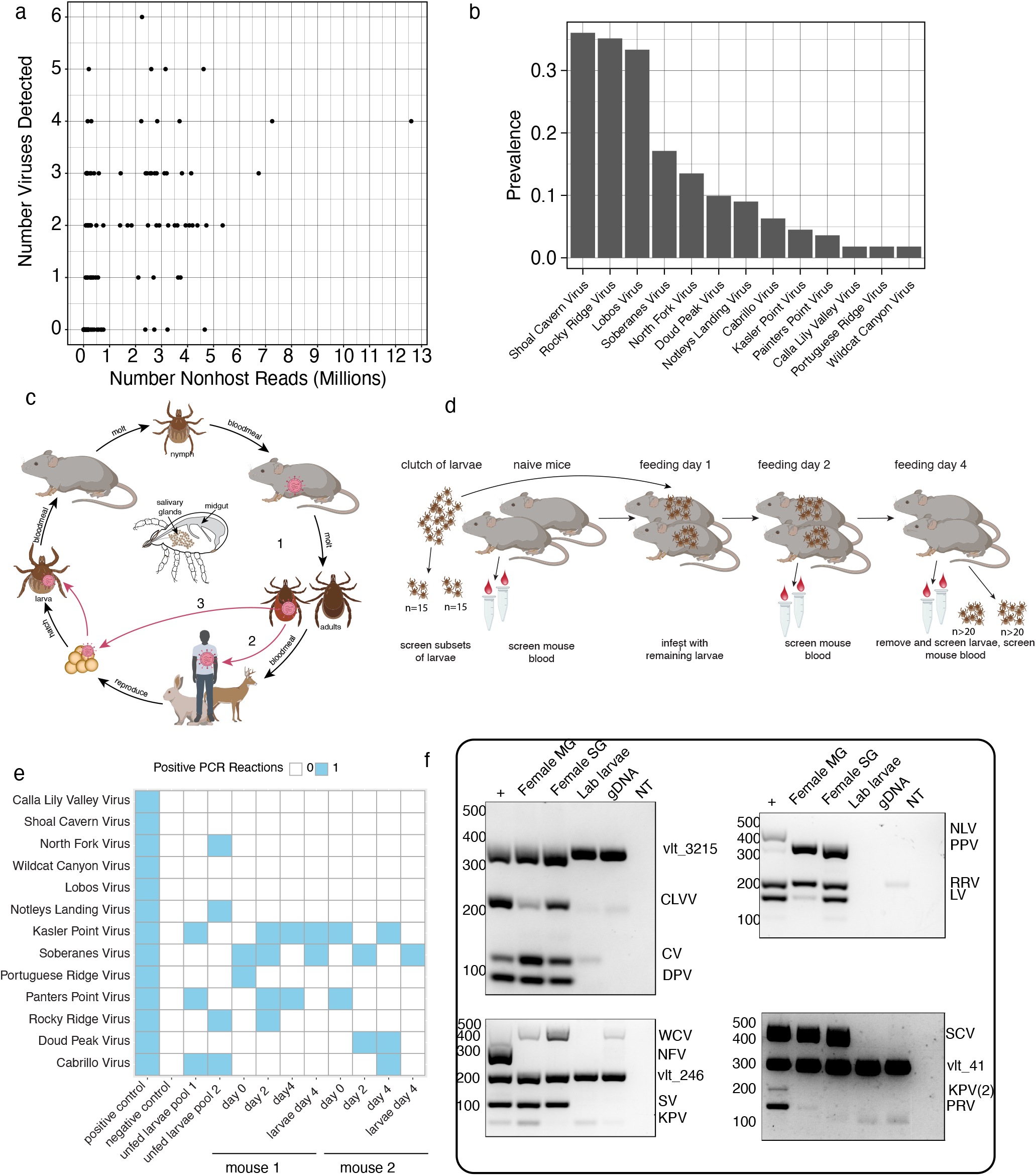
Discovered viruses in *I. pacificus*. a) Scatterplot of nonhost library size and number of viruses detected per sample. b) Prevalence of each viruse across the dataset. c) Schematic of I pacificus three stage life cycle. Viruses can be transmitted horizontally between hosts and ticks (1 and 2) or vertically from adult female to offspring (3). d) Schematic of mouse transmission experiment. A sample of larvae were screened for viruses as a pool prior to infestation. Remaining larave were used to infest two mice. Mouse blood was collected and screened prior to infestation, as well as on days 2 and 4 of feeding. On day 4 ticks were removed and screened as a pool. e) Summary of PCR results from experiment descrived in d. Unfed larvae pools are the same as in Figure 3d. f) Representative gel image of viruses tested in midguts and salivary glands and lab-reared larvae. Virus abbreviations are aligned to their expected band size. Bands beginning with “vlt” represent virus-like transcripts (see Figure 4).

**Figure S6:**
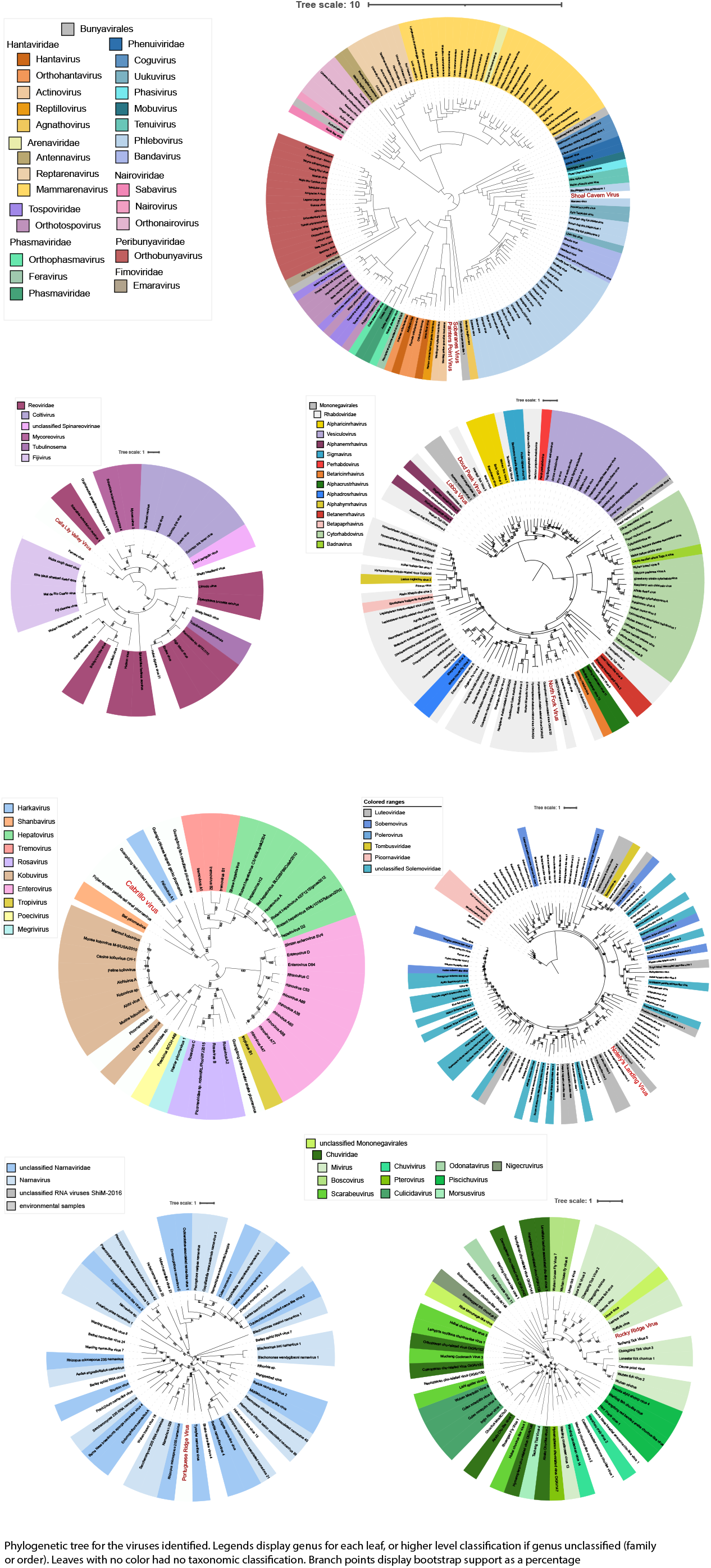
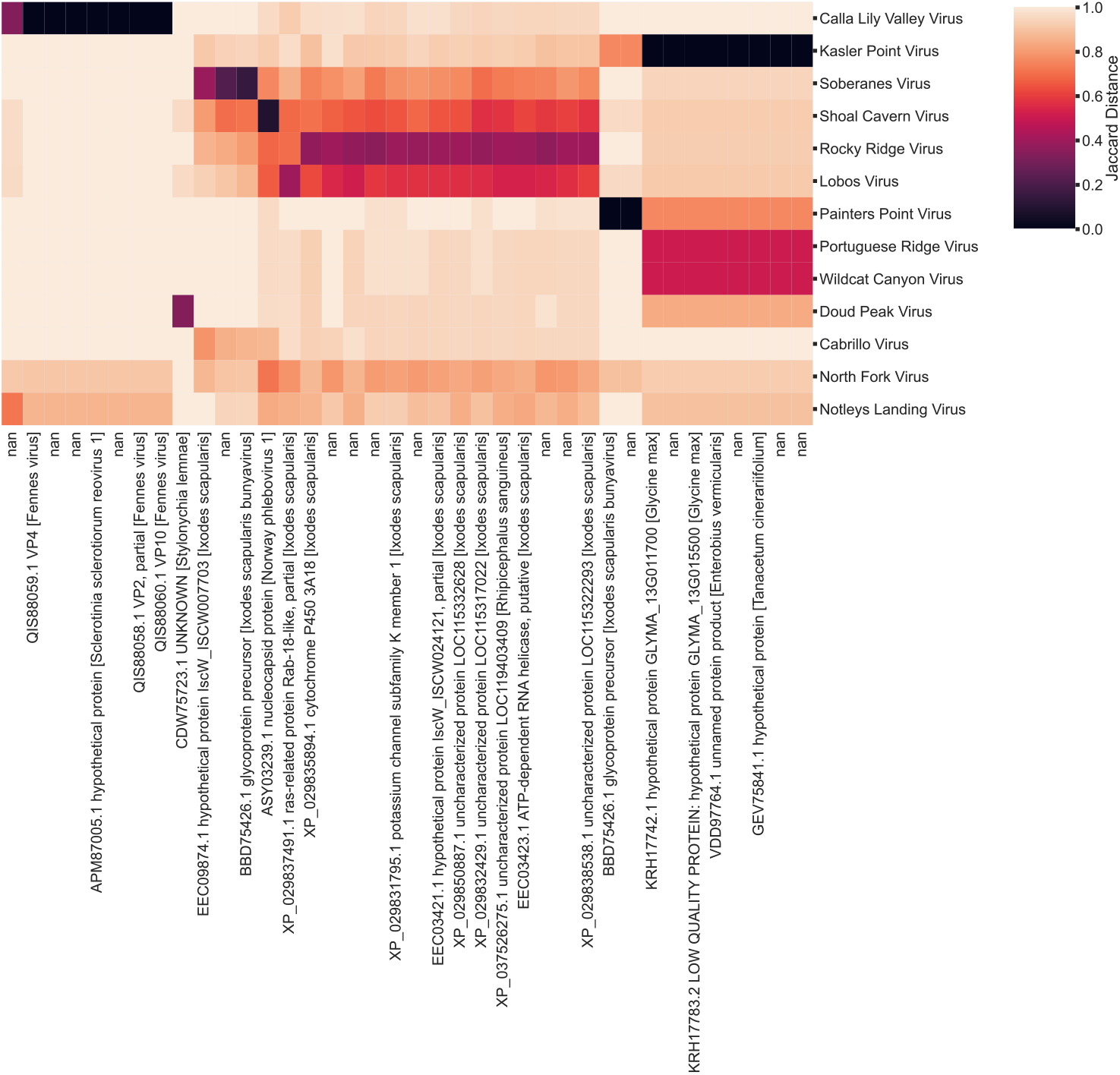
Identification of additional viral genomic segments. Heatmap of Jaccard distance of rdrp-containing contigs (rows) and other contigs in the dataset (columns). Columns are annotated with the closest blastx hit or nan if no hits were found.

**Figure S7:**
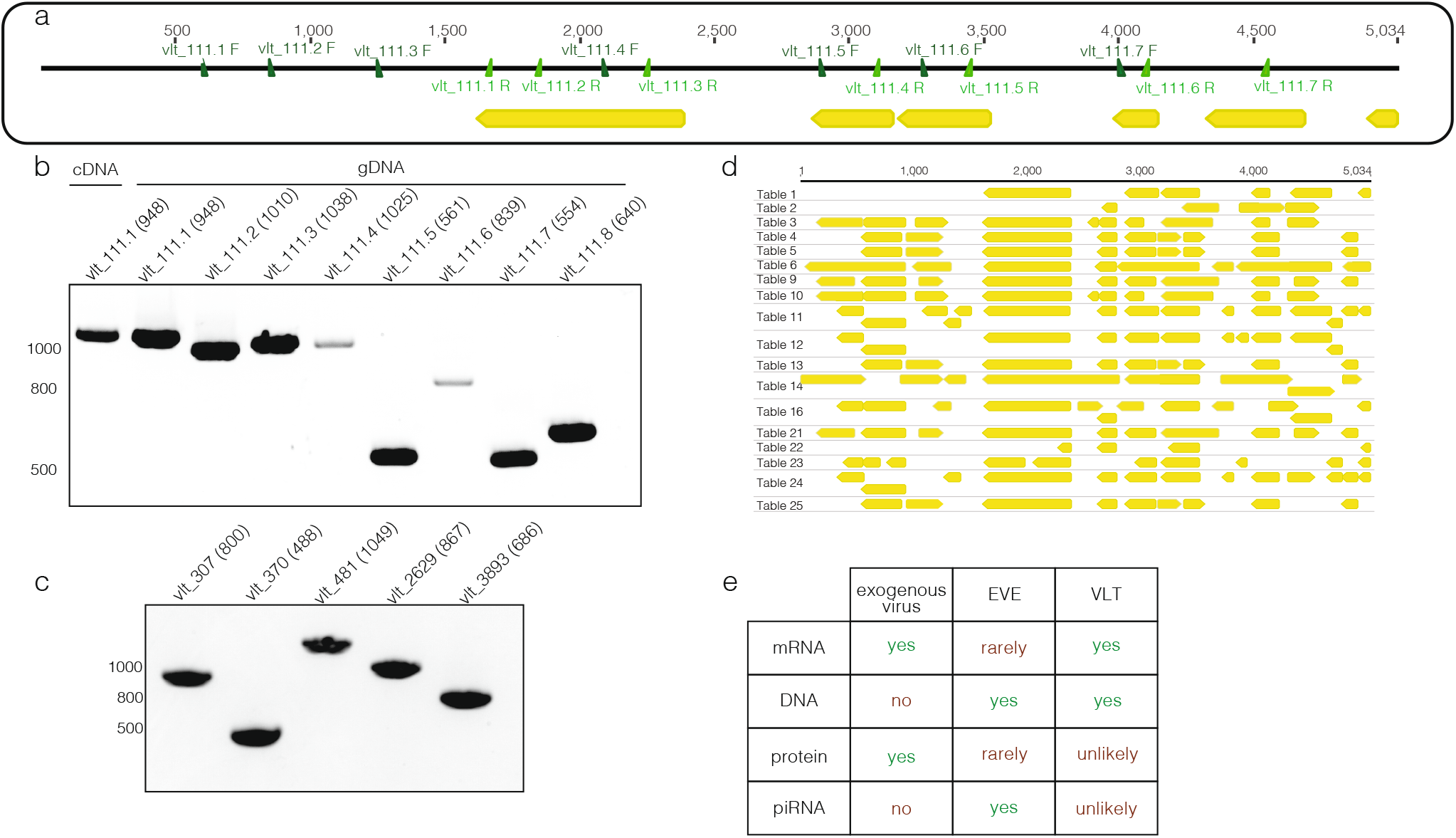
Confirmation of VLT sequence and presence in DNA. a)Visual representation of c111 identified from RNA seq, including predicted open reading frames (yellow) and primer pairs b) PCR reactions amplifying the regions indicated in a, nucleic acid type indicated on lane. Expected band size in base pairs indicated in parentheses. c) PCR reactions from an additional 5 virus-like sequences amplified from gDNA, expected band size indicated in parentheses. d) Open reading frames predicted for c111 using alternative codon tables e) Table summarizing expected pattern of exogenous viruses, endogenous viral elements (EVEs) and observed pattern of VLTs

**Figure S8:**
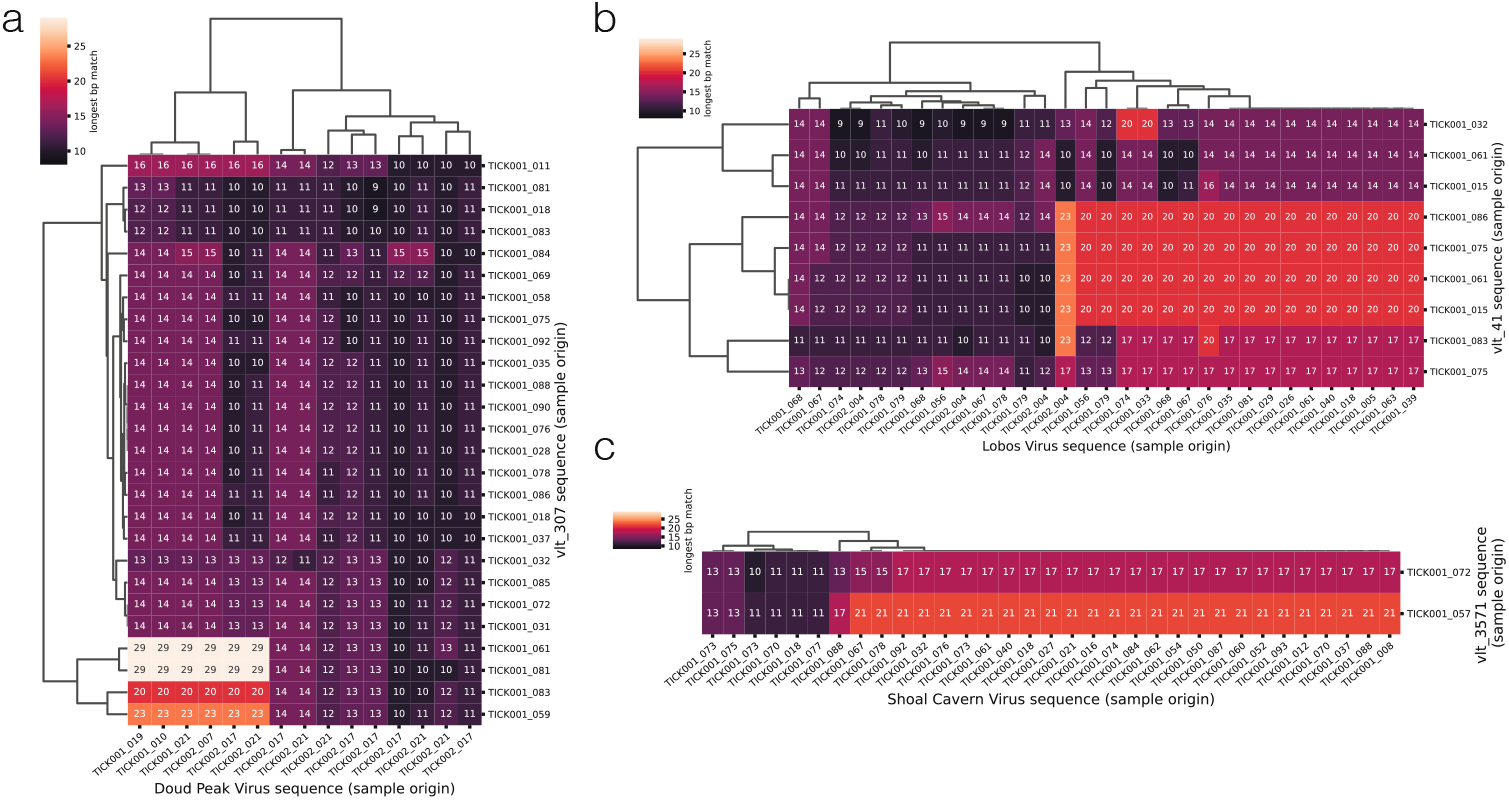
Comparision of individual VLT to virus sequences. a) Clustermap displaying the length of the longest perfectly matching sequence between each sequence assigned to the *Doud Peak Virus* cluster and each sequence assigned to the vlt_307 cluster. Rows and columns are labeled with the tick sample from which the sequence originated b-c) As in a but for *Lobos Virus*:vlt_41 and *Shoal Cavern Virus*:vlt_3571 respectively

